## Supplementary material for "Cardio PyMEA: A user-friendly, open-source Python application for cardiomyocyte microelectrode array analysis": S1 File - Tutorial

**Tutorial for adding new MEA geometries to Cardio PyMEA**

This tutorial will explain how to add new MEA geometric configurations (i.e. MEAs of different inter-electrode spacing and electrode diameters) to Cardio PyMEA.

In Cardio PyMEA, MEA geometries are stored in dictionaries. Dictionaries are a particular type of variable in Python. Each dictionary contains a {key: value} pairing and is denoted by curly braces (or, alternatively, can be represented as: dict(key: value). These dictionaries are used by the software to assign column labels (which, except for the first column, correspond to the electrodes in the raw data *.txt file) in the dataframe. This is how the coordinate system is utilized in the software: each electrode (column label) is referenced to the corresponding dictionary key to obtain its coordinates and generate the heat map.

The dictionaries used to assign MEA coordinates in Cardio PyMEA are housed in a class called ElectrodeConfig. This class primarily acts as a container for different MEA geometries. Each MEA geometry is contained in a dictionary assigned to a variable, e.g. *self*.mea_120_coordinates = {key: value} pairs, as shown in the screenshot T1 Fig below. If you are new to programming, we will pause here briefly to explain the variable naming scheme. If you are not new to programming and understand the naming scheme of the previous variable, please move to the next paragraph. The variable *self*.mea_120_coordinates is referred to as a class ‘attribute’. Class attributes are denoted via dot notation following the object the class refers to. In this case, the use of “*self*” describes the variable you named to instantiate the class (i.e. to initialize, or assign the class to a variable so that it can be used). For example, the class ElectrodeConfig is used to define the variable electrode_config = ElectrodeConfig(). This instantiates the class and assigns it to the variable *electrode_config*, which now takes on all of the properties (and attributes) of the class.

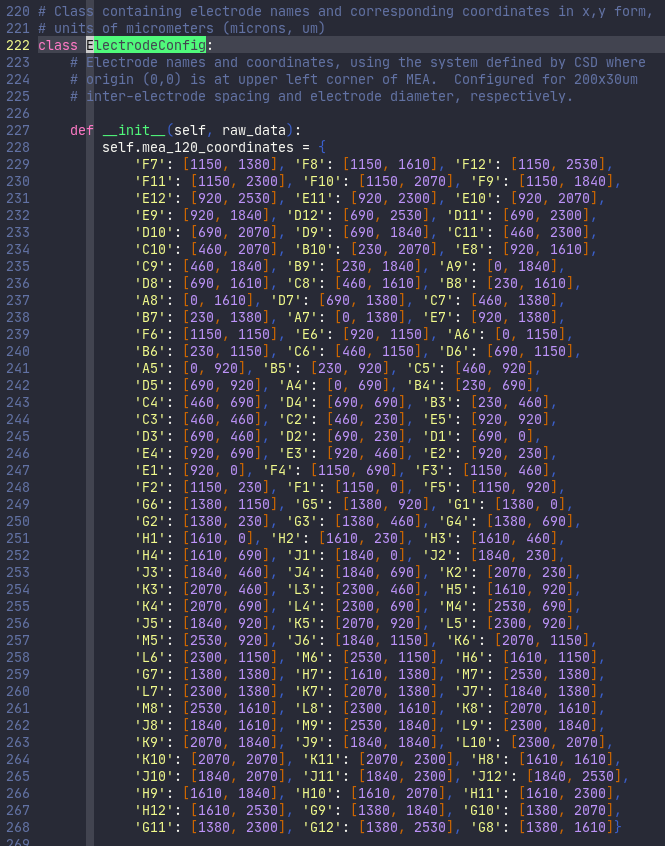

**T1 Fig. Screenshot of ElectrodeConfig class, showing one MEA dictionary.**

Consequently, the word “*self*” is, in this case, synonymous with “*electrode_config*”. Using the word “*self*” in the class allows the class to remain highly generalized to whatever variable name you later choose to instantiate it with. In other words, what you read as “*self*.mea_120_coordinates” in the above code will be referenced within the code as “*electrode_config*.mea_120_coordinates”, because that is how we chose to instantiate the ElectrodeConfig class (i.e. by declaring the variable electrode_config = ElectrodeConfig()).

In order to add a new electrode geometry to Cardio PyMEA, simply designate a new dictionary in the ElectrodeConfig class with the characteristics you desire. For instance, if you are using a 10 micron diameter, 100 micron interelectrode spacing, 120 electrode MEA, you could define your variable in the following way:

*self*.mea_120_10_100 = {key: value pairs}

Then, you would add your MEA labels, or names, as the keys in the form of strings (denoted by the use of quotation marks, “”), and the coordinates as the value pairs in the form of a list (denoted by the use of square brackets, []). You will want to make sure you follow your manufacturer’s schematic/order for the electrode labeling. This is to ensure that the first electrode column corresponds to the first electrode channel, as designated in the data file and manufacturer recording software. Please see the following link at Multichannel Systems website for additional information regarding electrode schematics: https://insert_the_correct_link_here.com.

Once you have finished setting up your MEA dictionary, you will next look at the electrode_toggle(self, raw_data) method (the term for a function that is housed within a class) in the ElectrodeConfig class, as shown in T2 Fig.

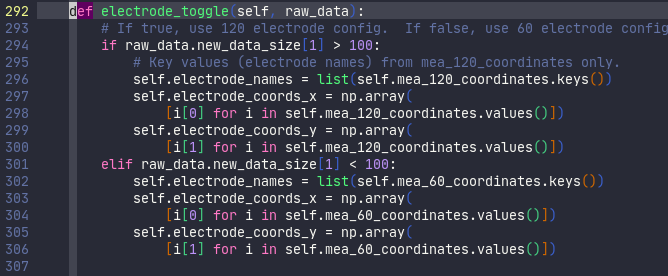

**T2 Fig. Screenshot of electrode_toggle method within the ElectrodeConfig class.**

This method is called during the data import stage (by the data_import function) to pick the correct MEA electrode labels and coordinates for application to the data file. In order to complete the implementation of your newly added MEA geometry, you need to update the conditional statements within this electrode_toggle method. As of 3/7/2022, Cardio PyMEA is set up to handle the simple case of determining whether an 60 or 120 electrode MEA is used, using admittedly crude decision-making criteria. Provided that you are also using either a 60 or 120 electrode MEA, the easiest mechanism through which to incorporate new geometries (i.e. a geometry that is distinct from our default 200 μm inter-electrode spacing and 30 μm electrode diameter) is to replace *self*.mea_120_coordinates or *self*.mea_60_coordinates inside the if-elif (interpreted as “if this, do this; else if that, do that”) conditional with your new variable name. If you are using an MEA with a different number of electrodes, you could easily add another “elif raw_data.new_data_size[1] < ##: condition” and replicate the lines in the existing if-elif conditional statements, updating your new lines for your new MEA.

After you have completed all of the above, save the analysisGUI.py file and run the program from the terminal. It should launch and, provided there are no typographical errors and any added decision-making steps were implemented correctly, your new MEA geometry should be incorporated into the software. Congratulations!
